# Probing the stability of the SpCas9-DNA complex after cleavage

**DOI:** 10.1101/2021.08.04.455019

**Authors:** Pierre Aldag, Fabian Welzel, Leonhard Jakob, Andreas Schmidbauer, Marius Rutkauskas, Fergus Fettes, Dina Grohmann, Ralf Seidel

## Abstract

CRISPR-Cas9 is a ribonucleoprotein complex that sequence-specifically binds and cleaves double-stranded DNA. Wildtype Cas9 as well as its nickase and cleavage-incompetent mutants have been used in various biological techniques due to their versatility and programmable specificity. Cas9 has been shown to bind very stably to DNA even after cleavage of the individual DNA strands, inhibiting further turnovers and considerably slowing down in-vivo repair processes. This poses an obstacle in genome editing applications. Here, we employed single-molecule magnetic tweezers to investigate the binding stability of different *S. pyogenes* Cas9 variants after cleavage by challenging them with supercoiling. We find that different release mechanisms occur depending on which DNA strand is cleaved. After non-target strand cleavage, supercoils are immediately but slowly released by swiveling of the non-target strand around the DNA with friction. Consequently, Cas9 and its non-target strand nicking mutant stay stably bound to the DNA for many hours even at elevated torsional stress. After target-strand cleavage, supercoils are only removed after the collapse of the R-loop. We identified several states with different stabilities of the R-loop. Most importantly, we find that the post-cleavage state of Cas9 exhibits a higher stability compared to the pre-cleavage state. This suggests that Cas9 has evolved to remain tightly bound to its cut target.

## INTRODUCTION

CRISPR (Clustered Regularly Interspaced Palindromic Repeats)-Cas (CRISPR associated) systems constitute adaptive immune systems against nucleic acid-containing invaders in prokaryotic cells (1). CRISPR-Cas9 from the bacterial species *Streptococcus pyogenes* (SpCas9) is a 160 kD protein that forms a ribonucleoprotein (RNP) complex with CRISPR-RNA (crRNA) and trans-activating RNA (tracrRNA). This complex can site-specifically target double-stranded DNA (dsDNA) and induces a double-strand break using its RuvC and HNH nuclease domains (2–4). Since its discovery, SpCas9 and other Cas9 variants have been widely used in rapidly emerging genome engineering applications in prokaryotic, plant and animal cells (5–7). The target specificity is mainly encoded by the 20 nt spacer region at the 5’ end of the crRNA of the SpCas9 RNP complex. The tracrRNA and crRNA can be replaced by a single-guide RNA chimera (sgRNA) (4) which simplifies RNP formation. A prerequisite for site-specific DNA targeting is a three-nucleotide PAM sequence (protospacer-adjacent motif, NGG for SpSpas9) next to the target site. SpCas9 weakly interacts with suitable PAM sites via facilitated diffusion (8). PAM recognition triggers an initial melting of the DNA double strand and primes base pairing between the crRNA spacer and the DNA target strand (TS) while displacing the non-target DNA strand (NTS) such that a so-called R-loop structure is formed (9). R-loop expansion is initiated from the PAM proximal nucleotides over the entire spacer region and constitutes the actual recognition of matching targets. In case of strongly mismatched targets, R-loop expansion is impeded and stops or the R-loop collapses (10). Successful base pairing between TS and crRNA spacer is verified by two conformational changes of the SpCas9 RNP. Formation of an R-loop intermediate of 9 bp length triggers the transition into a “checkpoint” state, opening up a channel to further accommodate the TS-crRNA hybrid. This channel is formed by a conformational change of the REC2 and REC3 domains (11,12). In this state, the HNH domain is still positioned more than 30 Å away from the cleavage site in the TS. The second transition into the “docked” state is induced once the R-loop reaches the PAM-distal end, causing a large movement of the HNH domain to reach its site of catalytic activity (13–15). Mg^2+^ ions as cofactor and a maximum of three terminal mismatches are prerequisites for the transition into the docked state that allows DNA cleavage. The TS and NTS are cleaved by the HNH and the RuvC domains, respectively, followed by a further conformational change into a post-cleavage product state (16,12). In this state, the HNH domain undocks from the cleavage site and is thought to become completely disordered. After inducing a double-strand break, an extremely slow release of the DNA products essentially limits SpCas9 to undergo further turnovers (17). This results in an inhibited and slow *in-vivo* DNA repair process (18,19). However, even for target bound SpCas9, the 3’ end of the cleaved NTS can be released and targeted by complementary ssDNA or 3’ to 5’ ssDNA exonucleases (16,19).

Introducing inactivating point mutations at positions D10A (RuvC) or H840A (HNH) turns SpCas9 into TS or NTS nickases (4). Cas9 nickase mutants and cleavage-incompetent Cas9, which carries both mutations, have been used in various genome editing applications to increase specificity or to avoid double-strand breaks (20,21). Fusion of Cas9 variants with other enzymes, such as the repair enzyme Rad51, the endonuclease FokI, or an engineered reverse transcriptase, expanded the repertoire of Cas9-mediated functionalities (22,23,21). While a large body of research has investigated the structure of the conformational states of SpCas9 during R-loop formation and after DNA cleavage, the stability of the SpCas9-DNA complex in these states and their dependence on supercoiling remains elusive. Developing a thorough understanding of the post-cleavage stability of SpCas9 and the respective mutants can be helpful to improve the efficiencies of genome editing techniques.

Here, we investigated the post-cleavage state stability of SpCas9 and its respective nickase mutants by challenging it with supercoiling using single-molecule magnetic tweezers experiments (9,24,25). We show that SpCas9 bound to a target with a dsDNA break or a nicked NTS can easily relax superhelical tension. It can thus not be dissociated in processes that generate this kind of mechanical stress, such as transcription and replication (26–28). In contrast, SpCas9 bound to an intact target or a target with a nicked TS cannot release superhelical tension and can thus be dissociated from the DNA when elevated tension is induced. In both cases, protein states with different stabilities are found indicating dynamic structural transitions in the post-celavage state. We show that dissociation from a target DNA with a nicked TS requires considerably higher levels of supercoiling compared to an intact target, suggesting an increased stability of the post-cleavage conformational state.

## MATERIAL AND METHODS

### DNA substrates for magnetic tweezers measurements

Double-stranded DNA constructs for magnetic tweezers experiments with lengths of 1500 bp and 2700 bp were prepared as previously described (25). A single copy of a given protospacer matching the spacer of the SpCas9 sgRNA and including a 5’-AGG-3’PAM on the non-target strand (see Table S1) was cloned into the SmaI site of a pUC19 plasmid. From the plasmid, a 1.5 kbp or 2.7 kbp fragment including the SpCas9 target site was amplified by PCR, using primers in which either a NotI or a SpeI restriction enzyme site was introduced (see Table S1). After digestion with NotI and SpeI, the fragment was ligated at either end to ∼600-bp-PCR fragments containing multiple biotin (SpeI site) or digoxigenin (NotI site) modifications (29).

### sgRNA synthesis for magnetic tweezers experiments

The template for T7 transcription of the SpCas9 single sgRNA was generated by PCR, using the High Fidelity Phusion Polymerase (Thermo Fisher Scientific) and overlapping oligonucleotides. RNAs were produced by *in vitro* transcription using the TranscriptAid T7 High Yield Transcription Kit (Thermo Fisher Scientific) and purified using the GeneJet RNA Cleanup and Concentration Kit (Thermo Fisher Scientific). The sequences of the oligonucleotides and sgRNA used in this study are available in Table S1.

### Preparation of SpCas9

Plasmids for the expression of wildtype SpCas9 and the catalytically inactive SpCas9 variants carrying the mutations D10A/H840A were obtained from Addgene (plasmid ID: 39312 and 39318). SpCas9 variants that carried a single mutation of the catalytic site (D10A or H840A) were generated by site-directed mutagenesis using the wildtype SpCas9 expression plasmid as template. Expression was performed *in E. coli* BL21 (DE3) cells. Expression and cell lysis conditions were chosen as described by Jinek et al. (4). The cleared cell lysate was applied to a HisTrap FF Ni-NTA affinity column (GE healthcare) equilibrated with HisA buffer (20 mM HEPES pH 7.5, 100 mM NaCl, 5 mM MgCl_2_, 5% (v/v) glycerol, 20 mM imidazole). The column was washed with 2 column volumes (cv) of HisA buffer and 10 cv of HisA buffer containing 1 M NaCl to liberate unspecifically bound nucleic acids from SpCas9. Subsequently, the column was equilibrated with 2 cv of HisA buffer and pre-eluted with 2 cv buffer HisA with increased imidazole concentration of 40 mM. In the final elution step, the column was washed with 5 cv elution buffer (20 mM HEPES pH 7.5, 100 mM NaCl, 5 mM MgCl_2_, 5% (v/v) glycerol, 200 mM imidazole). Elution fractions were pooled and digested with 50 U AcTEV protease (Thermo Fisher Scientific) over night at 4 °C. The cleaved His6-MBP-tag was removed by gel filtration using a HiLoad 16/60 Superdex 200 column (GE healthcare) equilibrated in size exclusion buffer (20 mM Tris-HCl, 150 mM NaCl, 5 mM MgCl_2_, 5% (v/v) glycerol, 1 mM DTT). Elution fractions contained nucleic acid free SpCas9 or SpCas9 variants (A260/280 < 0.6). Fractions were flash frozen in liquid nitrogen and stored at −80 °C.

### Reconstitution of guideRNA-SpCas9 complexes

For characterizing DNA cleavage, SpCas9 was pre-loaded with either a sgRNA or a pre-annealed crRNA:tracrRNA complex (Table S1 and S2). To reconstitute SpCas9-RNA complexes, a stock solution containing 1 µM SpCas9 and 1 µM guide RNA in 1x Cas Buffer (20 mM Tris–HCl pH 7.5, 100 mM NaCl, 10 mM MgCl_2_, 2% (v/v) glycerol, 1 mM DTT, 0.05% (v/v) Tween20) was incubated for 10 minutes at room temperature followed by a centrifugation step (20.000 g, 5 min, room temperature).

### Plasmid-DNA cleavage assays

As substrate for plasmid-DNA cleavage assays, we cloned target plasmid (pM53.1; 6290 bp) containing the sequence of the *S. cerevisiae* PHO5 gene with its native genomic context 500 bp upstream and downstream of the gene in a pBluescript vector backbone. The PHO5 locus was amplified from yeast genomic DNA using primers 5’-GGCCGCAGATCAAGTCAGAG-3’ and 5’-GCAAGTCACGAGAAATACCA-3’. The gene is flanked by a *Z. rouxii* sequence-specific recombinase R recognition site (RS) upstream, and a triple *E. coli* LexA binding site downstream of the gene followed by another RS element. The plasmid was generated making use of pM49.2 containing the upstream RS element and the downstream triple LexA binding site and RS element (for details of the cloning procedure see (30)). Reconstituted SpCas9-sgRNA complexes were added in 10-fold excess to 5 nM pM53.1 plasmid DNA containing the target sequence and a 5’-AGG-3’ PAM motif in 1x Cas buffer. For kinetic analysis, 15 µl samples were taken at different time points and the reactions were stopped by addition of EDTA (83 mM final concentration). Subsequently, SpCas9 was digested by the addition of 0.36 U Proteinase K (Thermo Fisher Scientific) followed by an incubation at 55 °C for one hour. After Proteinase K digestion, 6x DNA loading dye (10 mM Tris-HCl pH 7.5, 0.03% bromophenol blue (w/v), 60% glycerol (v/v), 6 mM EDTA) was added to the samples and fragments were resolved on a 0.9% agarose gel. Cleavage products were visualized using a ChemiDoc Imaging System (Bio-Rad) after ethidium bromide staining of the agarose gel.

### Short target DNA cleavage assay for determination of SpCas9 strand-specific cleavage

Fluorescently labeled double-stranded target DNA was formed annealing two single-stranded oligonucleotides. Single strands were purchased from Microsynth Seqlab (Göttingen, Germany) with a fluorescent label coupled to the 5’-end. The target strand carried a Cy3 label and the non-target strand a Cy5 label (Table S2). Annealing was performed by incubation of equimolar amounts of TS and NTS in annealing buffer (final concentration: 3 mM HEPES pH 7.4, 10 mM CH_3_COOK, 0.2 mM Mg(CH_3_COO)_2_ at 95 °C for 3 min and passive cool-down to room temperature. The reconstituted SpCas9-crRNA:tracrRNA complex (final concentration of 250 nM) was added to 5 nM of the double-stranded target DNA in a total volume of 10 µl in 1x Cas buffer. Cleavage reactions were incubated at 37 °C for 1 h and the reaction was stopped by the addition of 0.36 U Proteinase K (Thermo Fisher Scientific), followed by incubation at 55 °C for 45 min. The samples were mixed with loading dye (final concentration: 47.5% (v/v) formamide, 0.01% (w/v) SDS, 0.01% (w/v) bromophenol blue, 0.005% (w/v) xylene cyanol, 0.5 mM EDTA). Immediately prior to loading of the samples, the samples were denatured at 95 °C for 5 min. Denatured samples were separated for 30 min on a pre-heated 15% PAA, 6 M urea, 1× TBE gel (300 V, 45 min). Cleavage products were visualized using a ChemiDoc Imaging System (Bio-Rad).

### Magnetic Tweezers experiments

The measurements were performed in a custom-built magnetic tweezers setup allowing the GPU-assisted real-time tracking of the DNA length for up to 100 DNA molecules in parallel (31). DNA constructs for the tweezers experiments were bound to magnetic beads of 1 µm-diameter (MyOne; Invitrogen) and flushed into the glass flowcell of the setup allowing the anchoring of the digoxigen-modified end to the anti-digoxigenin coated surface of the cell (29,32). After removing unbound beads by flushing, force was applied by lowering the magnets towards the flow cell and DNA tethered beads were selected. Using bead tracking with a camera (Mikrotron EoSens), the length of the individual DNA molecules was determined and the applied forces were calibrated (33). During experiments, desired forces on the DNA construct could be set by placing the magnets at a particular distance from the flow cell according to the calibration results. Supercoiling of DNA was achieved by turning the magnets. The resulting torque that depended mainly on the stretching force was calculated based on previous theoretical work (34,35). Time trajectories of the DNA length were recorded at 120 Hz and smoothed with a sliding average to 6 Hz for analysis. For measurements with SpCas9, reconstituted SpCas9-sgRNA was added in tweezers buffer (20 mM Tris-HCl (pH 8.0), 100 mM NaCl, 10 mM MgCl_2_, and 1 mg/ml BSA) at a concentration of 0.25 nM or 0.5 nM. After adding SpCas9, DNA length changes were monitored in real-time. For experiments where we exclusively challenged the DNA-bound SpCas9 complex, SpCas9 was allowed to prebind to negatively supercoiled DNA in the flowcell for 20 minutes to ensure efficient R-loop formation on most molecules before the experiment was started. The successful formation of an R-loop was confirmed by the occurrence of a pronounced change in DNA length.

### Data Analysis

Data analysis for the tweezers experiments was performed with Origin (OriginLab) and custom-built Matlab (The Mathworks Inc.) scripts.

#### Supercoil release velocities

For Cas9_dHNH_ experiments, a slow supercoil release was observed. The velocities of the release were determined in Matlab by performing linear fits to the individual release events (see Fig. 2). The velocities at each particular torque were then plotted in a histogram and fitted with a Gamma distribution. The Gamma distribution describes the total waiting time *t* of *n* successive events with exponentially distributed dwell times with mean value *λ*. The Gamma distribution is given by:

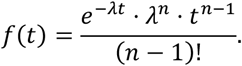

By using this distribution, we assume that SpCas9 releases the non-target strand one turn at a time, with each turn released after an exponentially distributed time with the same mean waiting time 1/*λ*.

#### R-loop collapse times

To determine transition times, R-loop transitions into the intermediate state or full collapse were fitted by step functions in Matlab. Survival plots of the collapse times were then fitted with bi- or tri-exponential functions using Origin.

## RESULTS

### Investigating the post-cleavage behavior of SpCas9 in single-molecule multiplex magnetic tweezers measurements

To investigate the post-cleavage behavior of different SpCas9 variants, we employed a single-molecule DNA twisting assay on a high-throughput magnetic tweezers platform (31). Numerous 2.7 kbp dsDNA molecules containing a single SpCas9 target site matching the crRNA spacer and multiple biotin or digoxigenin modifications at either end were attached on the respective DNA end to 1 µm streptavidin-coated superparamagnetic beads and to the antidigoxigenin-coated surface of the flowcell of the setup. The three-dimensional positions of multiple DNA-tethered beads were tracked in parallel in a home-built microscope (36) allowing the real-time observation of the DNA length of the molecules (Fig. 1A). The field gradient from a pair of magnets enabled the application of force onto the DNA molecule. The force could be controlled by adjusting the distance between the magnets and the flow cell. Turning the magnets allowed supercoiling of the DNA molecules in parallel.

**Figure 1.**
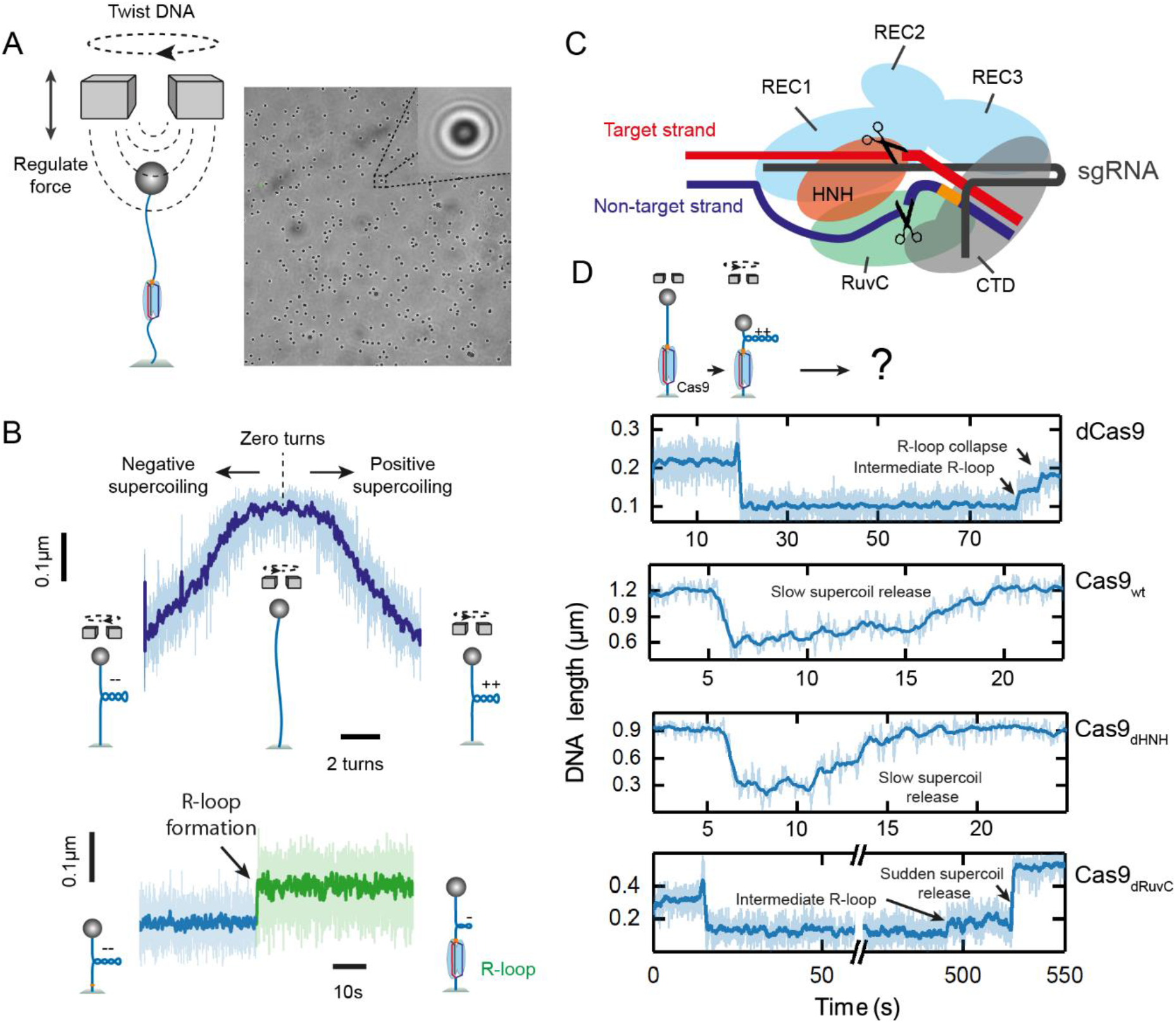
(A) Schematic representation of the magnetic tweezers assay used to twist single double-stranded DNA molecules that are tethered at either end to the surface of the fluidic cell and to a magnetic microbead. A pair of permanent magnets allows the application of a stretching force and molecule twisting. Up to one hundred beads were tracked in parallel (right: full microscope image with enlarged view of a single bead in the inset). (B) Top: Representative DNA supercoiling curve taken at 0.3 pN showing the characteristic DNA length reduction due to writhe formation upon twisting. Bottom: Representative time trajectory of the DNA length of a negatively supercoiled molecule (−6 turns, 0.3 pN) exhibiting a DNA length jump due to R-loop formation by Cas9_wt_. (C) Schematic representation of the SpCas9-sgRNA ribonucleoprotein complex forming an R-loop on double-stranded DNA. The scissors indicate the active sites of the RuvC and HNH nuclease domains. The orange section on the NTS represents the PAM. (D) Representative time trajectories when challenging R-loops formed by dCas9, Cas9_wt_, Cas9_dHNH_, and Cas9_dRuvC_ with positive supercoiling.

To monitor R-loop formation by SpCas9, we applied negative supercoiling which reduces the DNA length due to writhe formation (Fig 1B, top). R-loop formation results in untwisting of the DNA by about two turns, which absorbs part of the applied supercoiling such that it becomes visible as a sudden increase in the DNA length (Fig. 1B bottom).

As was previously shown for cleavage-incompetent dCas9 and other CRISPR-Cas effector complexes, the application of positive twist on DNA can promote the collapse of the R-loop and dissociation of the complex (9,25,10). In a simplified view, the R-loop is wrung out with the opposing superhelicity. At positive supercoiling, R-loop collapse on intact target DNA is also observable as a sudden DNA length change that, however, occurs in two steps (Fig. 1D, first trajectory). The first step corresponds to the transition into the checkpoint state, corresponding to the formation of the partial R-loop, while the second step corresponds to the total collapse of the R-loop and typically the dissociation of the protein.

We started to test the stability of the post-cleavage product state by challenging R-loops formed by wildtype SpCas9 (Cas9_wt_ thereafter) (Fig. 1C), SpCas9 with a mutated HNH domain (Cas9_dHNH_ thereafter), and SpCas9 with a mutated RuvC domain (Cas9_dRuvC_ thereafter) with positive supercoiling. Even though Cas9_wt_ induced double-strand breaks (DSBs) within ∼2 min after enzyme addition (Fig. S1), the DNA tethers in the tweezers stayed in place (Fig. S2) for several hours, indicating that Cas9_wt_ remained stably bound to both DNA ends as previously reported (13,16). We observed different behaviors of post-cleavage SpCas9-DNA complexes depending on which strand was cleaved (Fig. 1D). For Cas9_wt_ we did not observe the characteristic R-loop collapse with a 2-step supercoil release limited to ∼2 turns. Instead, an immediately starting but slow release of all applied supercoiling was observed (Fig. 1D, second trajectory). This behavior resembled that of eukaryotic topoisomerase IB, which nicks DNA and releases twist by a free swivel mechanism in which the release velocity is reduced by protein friction (37). Cas9_dHNH_, which solely nicks the NTS of the DNA, showed a similar behavior as Cas9_wt_ indicating that also in this case a swivel mechanism causes the slow release of supercoiling (Fig. 1D, third trajectory). In contrast, for Cas9_dRuvC_, which induces a nick on the TS, no immediately starting supercoil release was observed. Instead, the supercoiling level remained constant for prolonged times and was completely released in a fast, sudden step (Fig. 1D, fourth trajectory). Such fast supercoil release is indicative of a DNA nick in absence of bound protein (38) suggesting that Cas9_dRuvC_ suddenly dissociated leaving a free nick behind. This idea was supported by the observation that the fast release of all supercoils was frequently preceeded by a short substep indicative of the first step of R-loop collapse as seen for dCas9 (see small step in Fig. 1D, fourth trajectory).

### Bound Cas9_dHNH_ releases supercoiling through detachment of the NTS in a swivel-like manner governed by torque and friction

Next, we investigated the different responses of the SpCas9 variants to supercoiling after DNA cleavage in more detail. Both, Cas9_wt_ and Cas9_dHNH_ released supercoiling slowly by a swivel mechanism (Fig. 2A). The slow supercoil release was observed repeatably, indicating that the protein remained stably bound to the target. Otherwise, supercoils would be very rapidly released by a protein-free nick (38) and the introduction of additional supercoils in the DNA would be inhibited (as observed for Cas9_dRuvC_, see below). Hence, we observed a similar behavior for the Cas9_wt_ and Cas9_dHNH_ variants, which both nick the NTS, suggesting that occasional detachment of the NTS from the protein enables the swiveling/supercoil release (Fig. 2B). Despite the presence of a TS nick for Cas9_wt_, the DNA tethers did not rupture during supercoil release, indicating that the TS is tightly bound.

**Figure 2.**
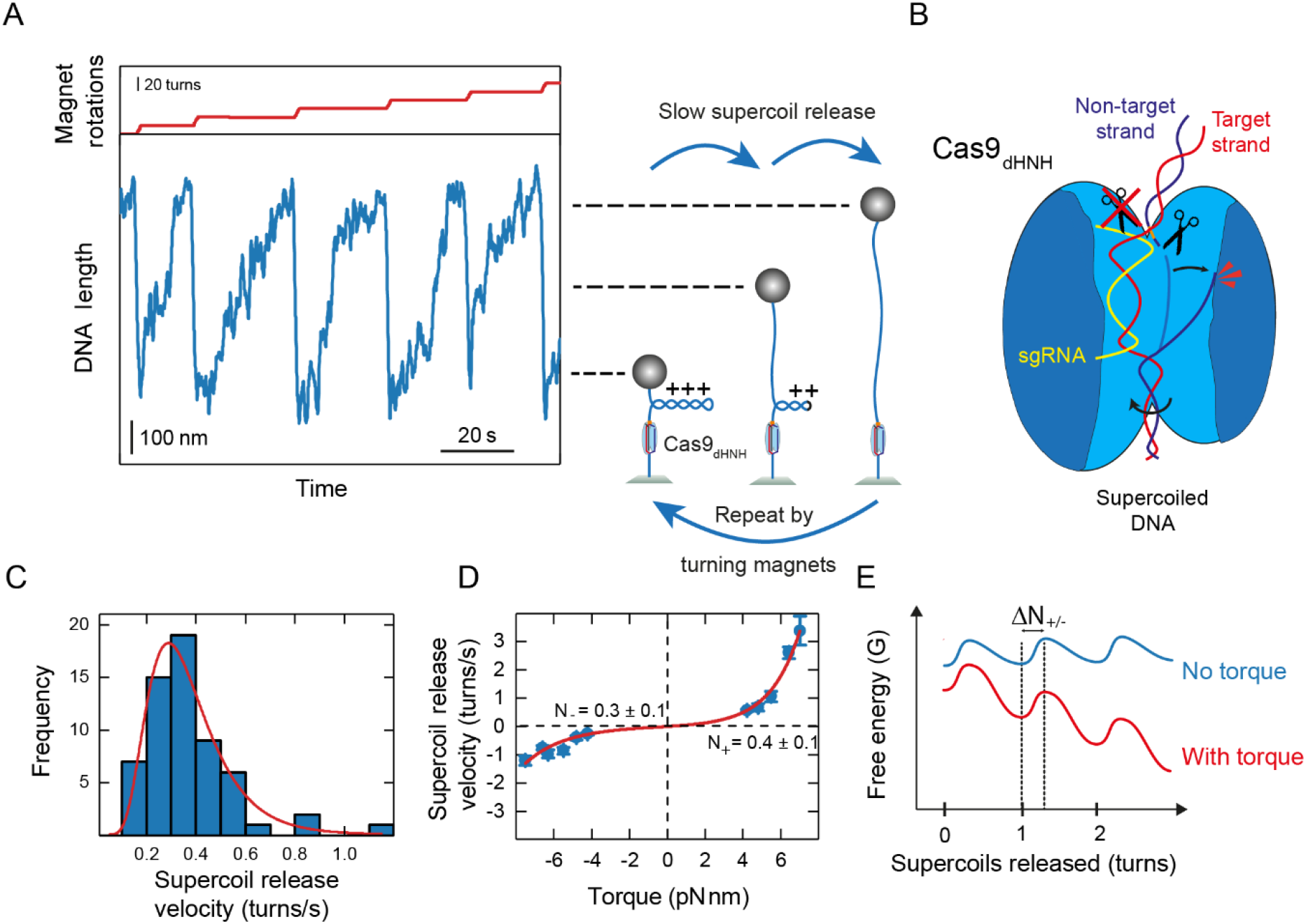
(A) Representative time trajectories of repeated supercoil release measured with magnetic tweezers (smoothed to 6Hz). Upon repeated step-wise DNA supercoiling, the DNA length is shortened. Supercoil release can be detected as a slow increase in DNA length, which is limited by friction due to the presence of Cas9_dHNH_. (B) Cartoon illustrating the supercoil release for DNA nicked by Cas9_dHNH_. When the NTS of a supercoiled dsDNA is cleaved, the applied torque is released by swivelling of the NTS around the TS. The swivelling is sterically obstructed by the protein. (C) Histogram of the measured supercoil release velocities given in turns per second (blue bars). The red line shows a fit with a Gamma distribution. (D) Torque-dependence of the supercoil release velocity (blue). The red line shows a fit of the data with Eq. (1) (E) Hypothetic scheme of the energy landscape for NTS release. The energy landscape is tilted downhill if positive or negative torque is applied (red line) in contrast to the absence of torque (blue line).

For Cas9_dHNH_, we carefully characterized the supercoil release velocity as function of the applied positive and negative supercoiling levels. For both supercoiling directions, a similar release behavior was recorded. Application of different forces allowed varying the acting torque during supercoil release. We measured the velocity of individual release events by applying a linear fit through each event. At each respective torque, we determined the linear slopes of the supercoiling curve which allowed us to convert the velocity from nanometers per second to turns per second. The distributions of the release velocities had a pronounced maximum and were well described by a Gamma distribution (Fig. 2C). The Gamma distribution describes a process containing *n* substeps with exponentially distributed dwell times (see Methods).

For the parameter which describes the number of substeps in the process, we obtained on average *n* = 6.6 ± 1.3 and *n* = 4.3 ± 0.3 for positive and negative supercoiling, respectively. These numbers agree well with the number of turns within the linear regime of the supercoiling curve, indicating the release of supercoils one turn at a time. The mean release velocities approximately showed an exponential increase with the applied torque for positive and negative supercoiling (Fig. 2D). Following Arrhenius transition-state rate theory, we modeled the supercoil release as a step-wise process in which supercoils can be released (or introduced) as single full turns. For this process, a periodic energy landscape can be drawn with energy minima at full turns (corresponding to the NTS docked onto Cas9_dHNH_) and transition state maxima in between (corresponding to a detached NTS and partially turned DNA) (Fig. 2E). Applied torque tilts the energy landscape to favor supercoil release. The torque dependence can be described using an exponential Arrhenius-term in which the activation energy required to reach the transition state is changed by the work to reach the transition state in presence of the torque. To consider supercoil release in positive and negative direction and to ensure a zero net release at zero torque, two torque-dependent Arrhenius-terms were used:

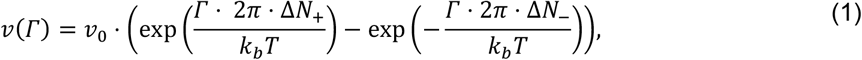

where *v*_0_ is the spontaneous swiveling rate in either direction at zero torque, *Γ* is the applied torque, and Δ*N*_+/−_ are the angular distances of the transition state from the energy minima for positive and negative supercoil release, respectively. Fitting this model to the data in Figure 2C resulted in transition state distances of Δ*N*_+_= 0.4 ± 0.1 turns and Δ*N*_−_= 0.3 ± 0.1, indicating that the transition state is located approximately half a turn from the energy minima. The swiveling rate at zero torque was *v*_0_= 0.04 ± 0.01 turns/s, i.e. Cas9_dHNH_ would spontaneously introduce or deplete a single supercoil every ∼25s even for relaxed DNA.

Overall, this data supports a model in which Cas9_dHNH_ releases the NTS by a swiveling mechanism governed by torque and friction. Due to the slight difference in the transition state distance, the release occurred slightly faster at positive than negative torque.

### TS cleavage causes two differently stable SpCas9 states

In a next step, we explored the stability of Cas9_dRuvC_ in the post-cleavage state on DNA when subjected to supercoiling. Cas9_dRuvC_ only cleaves the TS, such that supercoils cannot be released by NTS detachment and swiveling around the protein (Fig. 3A). Experimentally, this is seen as a constant supercoiling level over extended time periods (Fig. 3B). Full supercoil release can be observed as a sudden rapid step similar to the free nick generated by a nicking enzyme (38). This suggests that the twist is only released after R-loop collapse. Notably, the sudden supercoil release due to R-loop collapse is preceeded by a substep corresponding to a partial R-loop collapse i.e. the transition into the intermediate R-loop state. Subsequent repeated introduction of additional supercoiling failed, indicating that the enzyme dissociated upon R-loop collapse. After a waiting time of several minutes, the introduction of supercoils became possible again, indicating that a new Cas9_dRuvC_ molecule had bound to the nicked target site (Fig S3).

**Figure 3.**
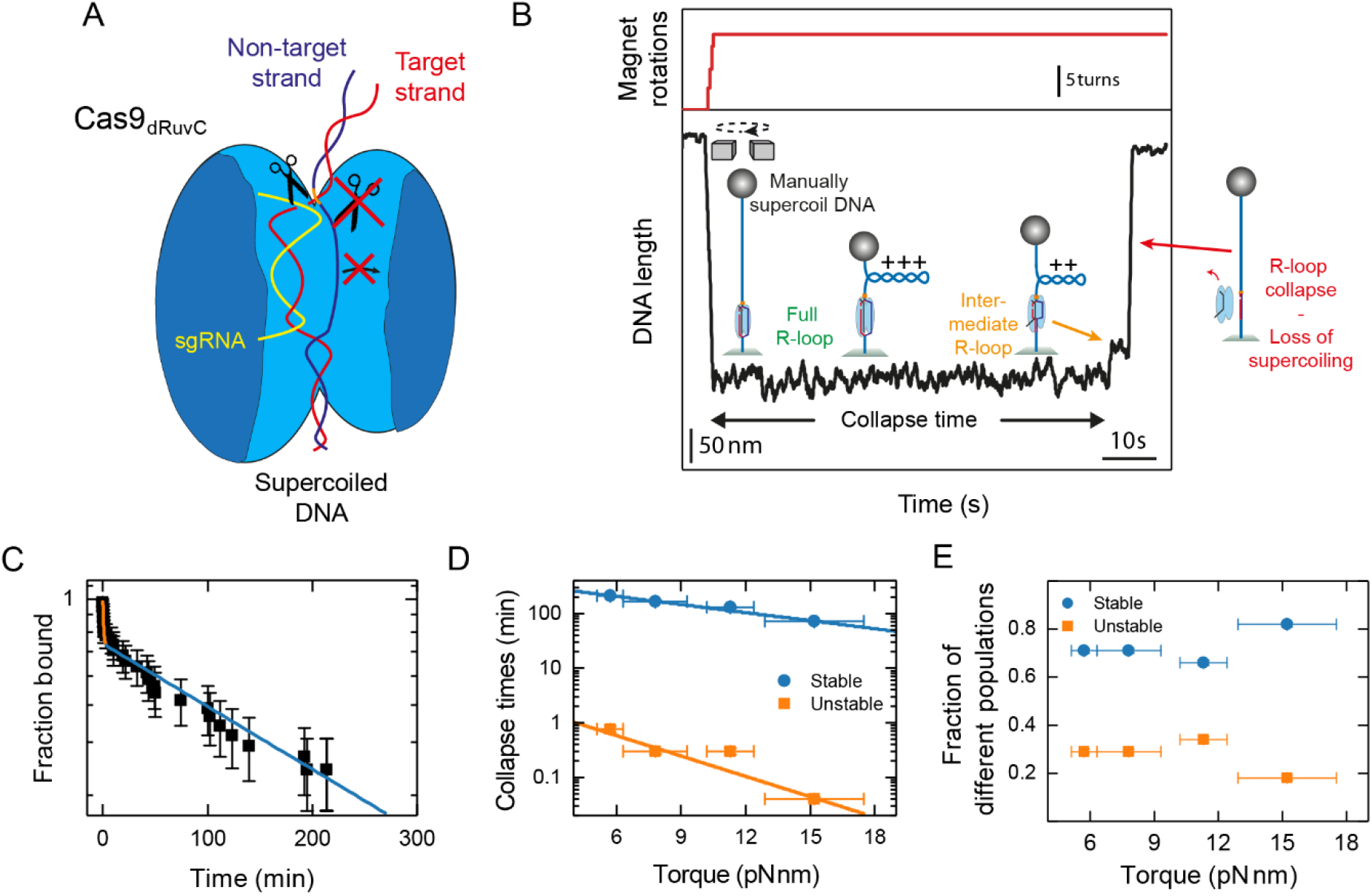
(A) Cartoon illustrating the nicking of the TS by Cas9_dRuvC_. The TS is nicked but remains tightly bound to Cas9_dRuvC_ such that supercoils are not released in presence of the protein. (B) Representative magnetic tweezers trajectory (smoothed to 6 Hz) showing data for an experiment in which Cas9_dRuvC_-nicked the DNA and the complex is challenged by strong positive supercoiling (high positive torque). No supercoils are released until part of the R-loop collapses (see orange arrow). This is followed by a rapid release of all supercoiling as observed for nicked DNA in absence of protein (red arrow). (C) Survival plot of the introduced R-loops (black squares) fitted with a bi-exponential decay function (orange/blue line). (D) Characteristic time constants for the R-loop collapse as function of torque obtained from the bi-exponential fits revealing an unstable (orange) and a stable (blue) population of R-loops. (E) Torque-dependence of the fractions of unstable (orange) and stable (blue) R-loops.

We carefully investigated the stability of Cas9_dRuvC_ bound to the nicked target site. To this end, we incubated the protein and DNA in the flowcell at negative supercoiling to allow R-loop formation. We then induced positive twist on the DNA to enforce the collapse of the R-loop and characterized the time between introduction of positive supercoiling and the collapse of the full R-loop. The collapse times were best described by a bi-exponential distribution (Fig. 3C). We, therefore, propose the existence of two distinct states distinguishable by their collapse times which were on the order of minutes for the unstable state and hours for the stable state. Repeating the experiment at different applied torques, we observed an exponential dependence of the R-loop collapse time on the torque for both states (Fig. 3D). This indicated the presence of a transition state for R-loop collapse (similar to the supercoil release above) as observed previously for various CRISPR-Cas proteins (25,10,39,9). Fitting the torque dependent data with a single Arrhenius function provided angular distances to the transition states of 0.18 ± 0.03 turns and 0.07 ± 0.01 turns for the unstable and stable state, respectively.

This corresponds to a DNA rewinding (most likely at the PAM-distal end) of about 2 and 1 bp. Furthermore, the fitting allowed to extrapolate the collapse times to zero torque for which we obtained 408 ± 58 min and 3.2 ± 1.5 min for the stable and the unstable state, respectively. On average, 72 ± 3% of the R-loops were in the stable state and 28% ± 3% in the unstable state with no obvious torque-dependence (Fig. 3E). We did not observe any dsDNA release by Cas9_wt_ until many hours after cleavage. This suggests that the occurrence of the unstable state after TS cleavage is unique to Cas9_dRuvC_ and does not occur in case of Cas9_wt_, since stable TS capturing would only be possible for a formed R-loop.

Overall, the results support that Cas9_dRuvC_ stays tightly bound to the target after TS cleavage until the full R-loop collapses. During this time, the Cas9_dRuvC_-sgRNA-DNA complex can adopt different conformational states, possibly in a dynamic manner.

### The post-cleavage product state stabilizes the R-loop in a torque-dependent manner

To reveal the relative stability of the post-cleavage state, we finally investigated the stability of SpCas9 in the pre-cleavage state. To this end, we challenged intact R-loops formed by a catalytically fully inactive dCas9 variant with positive supercoiling (Fig. 4A). The measurements were performed in presence of magnesium, allowing dCas9 to adopt the “docked” conformation of the HNH domain. As described before, R-loops formed by dCas9 collapsed in two steps at positive torque (Fig. 4B). The first step can be attributed to the transition from the docked to the intermediate state as observed for Cas9_dRuvC_. The second step, which was limited only to partial supercoiling release due to the lack of a DNA nick, can be attributed to the collapse of the intermediate state and dissociation of dCas9. Allowing a new R-loop formation at negative turns, the collapse process could be repeated multiple times (Fig. S5). The R-loop collapse times were best described by an exponential distribution with three distinct components, therafter called stable, semi-stable, and unstable state (Fig. 4C). As for Cas9_dRuvC_, R-loop collapse occurred in two distinct steps for all three components. The respective collapse times fell into the range of hours, minutes, and seconds with each component showing a slight torque dependence (Fig. 4D). Compared to Cas9_dRuvC_, the collapse time for the stable state was slightly decreased, while the collapse time of the unstable state did not significantly change. The occurrence of a torque-dependent semi-stable state was not observed for Cas9_dRuvC_. This may indicate flexibility of the docked state of dCas9. Similar to Cas9_dRuvC_, the fraction of molecules in the unstable state did not change with torque (Fig. 4E). However, the fraction of molecules in the semi-stable state increased, while the fraction of molecules in the stable state decreased with increasing torque. Due to the torque dependence of the fraction of molecules found in the different populations, the mean time of the R-loop collapse was strongly dependent on the torque applied (Fig. 4F). When comparing these results to Cas9_dRuvC_ (Fig. 4F), we found that the mean time of the R-loop collapse was generally higher for Cas9_dRuvC_ compared to dCas9. This indicates a further stabilization of SpCas9 after TS cleavage. The stabilization was found to be torque-dependent with pronounced stabilization at high torques. To correlate the two states with different stabilities to known conformations of SpCas9, we carried out experiments under Mg^2+^-free conditions at which the docked state of SpCas9 is suppressed (14). Even though the docked state is suppressed, a full R-loop is formed (Fig. S6A). We obtained an average collapse time that matches the collapse time of the previously observed unstable state (Fig. S6B). This suggests that the stable and semi-stable states represent the docked conformation of SpCas9.

**Figure 4.**
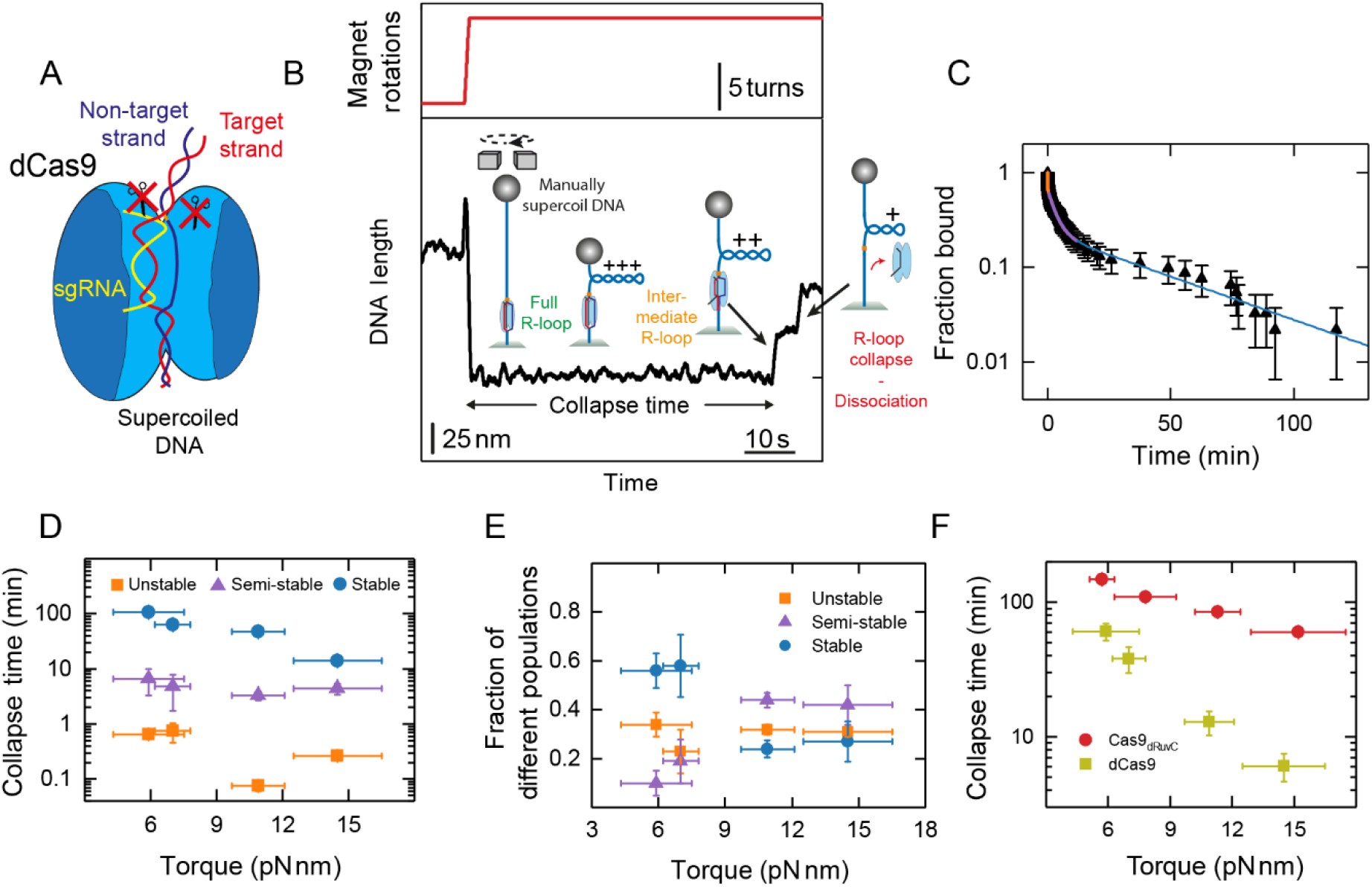
(A) Cartoon illustrating the pre-cleavage state of SpCas9, a state produced by the cleavage-incompetent dCas9 variant. (B) Representative magnetic tweezers trajectory (smoothed to 6 Hz) when challenging an R-loop formed by dCas9 with positive supercoiling. The R-loop spontaneously collapses in two characteristic steps. Absence of full supercoil release indicates that the DNA was not nicked by dCas9. (C) Survival plot of R-loops formed by dCas9 (black triangles) fitted with a tri-exponential decay function (orange/purple/blue line). (D) Characteristic time constants for the R-loop collapse as function of torque obtained from the tri-exponential fits revealing an unstable (orange), a semi-stable (purple), and a stable (blue) population of R-loops. (E) Torque-dependence of the fractions of unstable (orange), semi-stable (purple), and stable (blue) R-loops. (F) Mean R-loop collapse time averaged over all populations for dCas9 (green) and Cas9_dRuvC_ (red). Cleavage of the TS appears to stabilize SpCas9 on the DNA.

In conclusion, we showed that dCas9 adopts three different conformations when a full R-loop has been formed. Additionally, we observed a torque-dependent stabilization of the R-loop after cleavage of the TS.

## DISCUSSION

Several studies showed that SpCas9 stays tightly bound to its target DNA after cleavage of double-stranded DNA is completed. In this study, we elucidate the mechanisms by which SpCas9 achieves this high stability even under unfavorable conditions of positive supercoiling, a situation encountered during transcription and replication of genomic DNA. Using single-molecule magnetic tweezers measurements, we show that the post-cleavage behavior of SpCas9 strongly depends on which strand of the target site is cleaved.

NTS cleavage allows the NTS to detach from the protein and to swivel around the TS under friction. By this mechanism, even high supercoiling levels are quickly relaxed such that R-loop collapse and SpCas9 dissociation are circumvented. We determined a spontaneous swivel rate of 0.04 ± 0.01 turns/s at zero torque. This provides evidence for a spontaneous release of the NTS even in absence of supercoiling such that it becomes accessible for complementary ssDNA or exonucleases (16). Characterizing the torque dependence of this supercoil release allowed us to determine the transition state distances, indicating that the transition state lies at approximately half a turn.

A different behavior was observed for a SpCas9 variant that can bind the DNA target site but can only cleave the TS. Here, a facilitated supercoil release was impeded. As observed in the cryo-EM structure of the post-cleavage state of SpCas9 (12), the cleaved TS stays hybridized to the sgRNA in the post-cleavage state which can explain the absence of an immediate release of supercoiling. Applying elevated positive torque resulted in the collapse of the R-loop and dissociation of Cas9_dRuvC_. By carefully analyzing the collapse times, we observed two states of different stabilities with collapse times in the order of a few minutes and several hours, respectively. After TS cleavage, the HNH domain has been reported to be highly flexible (16,12). The different stability states observed in this work might stem from this flexibility. Additionally, there has been some debate on the dynamics of the pre-cleavage conformational states. While some studies have shown that SpCas9 reaches a stable, cleavage-competent ‘docking’ conformation in the presence of Mg^2+^ ions (14,40), other studies have observed only brief visits of the docked state with the majority of time spent in the ‘checkpoint’ conformation (15,41,42). Our results suggest that even though a complete R-loop is formed and cleavage is carried out, Cas9_dRuvC_ exhibits conformational flexibility, transitioning between an undocked state and a stabilized post-cleavage state. It is possible that the undocked state corresponds to the previously reported ‘checkpoint’ state. However, we note that even in the undocked state a full R-loop is formed, as confirmed by the extent of the length jump and the presence of the intermediate step during R-loop collapse (Figure S7). We show that the lifetime of the undocked state resembles that of a Mg^2+^-free control for which the “docked” state is not formed. For wildtype SpCas9, we did not observe DNA release after a cleavage of both DNA strands even after several hours of measurement indicating that it does not exhibit the undocked state after cleavage which was observed for Cas9_dRuvC_. Cleavage-incompetent dCas9 exhibits even three distinct states and, potentially, a higher conformational flexibility. Combining the results of Cas9_dRuvC_ and dCas9 we suggest that the stable state that is observed for both variants corresponds to the post-cleavage product state. This state would appear to be transiently visited even for the uncleaved target, albeit only at low levels of supercoiling. The semi-stable state is observed when dCas9 is bound to the target DNA without DNA cleavage. This state dominates under conditions of high supercoiling and most likely corresponds to the pre-cleavage state with a docked HNH domain. The unstable state is found for both dCas9 and Cas9_dRuvC_ and most likely represents the undocked state in agreement with the previously observed dynamic sampling between the different states (15,41,42) (Fig. 5). Overall, the dominance of the post-cleavage product state accompanying TS cleavage stabilizes the R-loop compared to the uncut target, especially at higher torques. This suggests that, under *in-vivo* conditions, SpCas9 has evolved to remain bound to its cut target. Possibly, this not only inactivates the invader DNA but also blocks transcription and/or replication of the foreign DNA, thereby fully blocking replication of a bacteriophage. However, this represents a challenge for efficient genome engineering applications.

**Figure 5.**
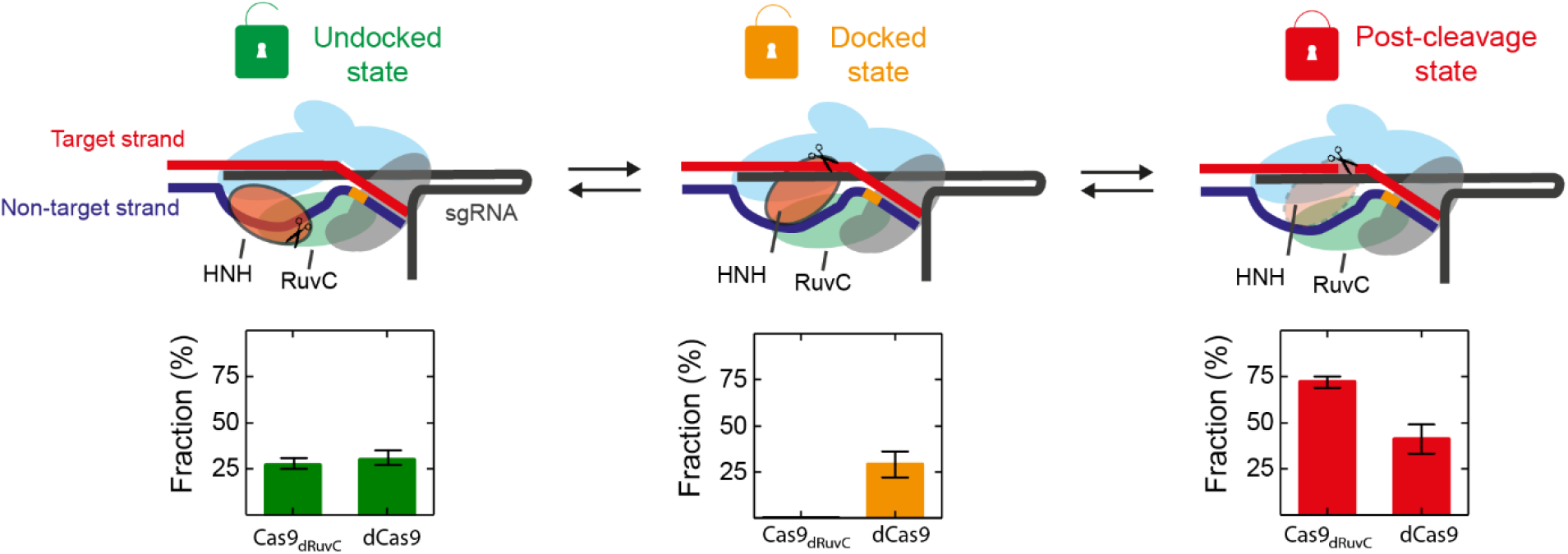
Schematic overview of the different states identified for SpCas9 bound to intact DNA and DNA with a nicked TS. We assigned the unstable state to the intermediate state with undocked HNH domain, the semi-stable state to the precleavage state with the HNH domain docked to the active site, and the stable state to the post-cleavage product state with a disordered (or flexible) HNH domain (12,16). Catalytically inactive dCas9 can occupy all states and most likely samples these states in a dynamic manner. Cas9_dRuvC_ was predominantly found in the stable (post-cleavage) state and rarely occupied the unstable state.

Overall, the presented results expand the knowledge about the post-cleavage state(s) of SpCas9 and the mechanisms by which SpCas9 avoids R-loop collapse and dissociation even under high twist that can be generated *in vivo* by the genome processing machineries. Nickase mutants of Cas9 have been used in various approaches for genome editing with strong differences in results depending on which nickase was used (22,43,44). Our findings may help to understand the described differences and might facilitate future genome editing applications with SpCas9 nickases.

## ACKNOWLEDGEMENT

We thank Joachim Griesenbeck for providing the pM53.1 plasmid. Furthermore, we thank Virginijus Siksnys, Tautvydas Karvelis, and Greta Bigelytė for providing the sgRNA.

