## Supplementary Information for "Probing the stability of the SpCas9-DNA complex after cleavage"

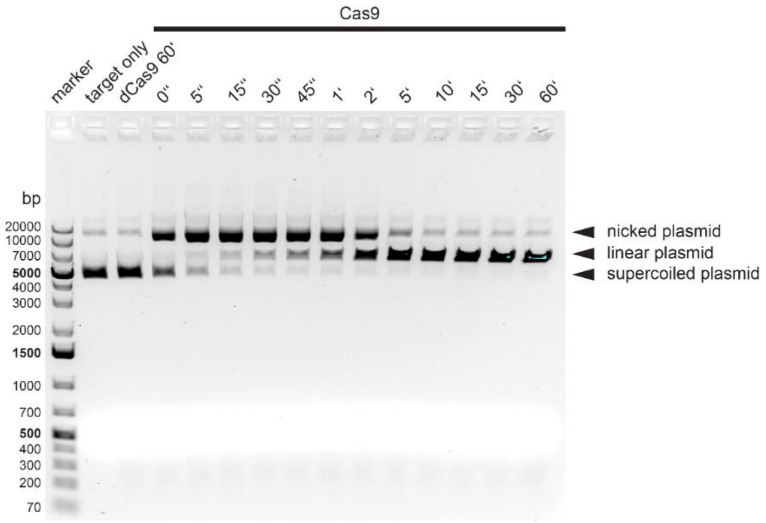

**Figure S1.** Plasmid DNA cleavage assay showing the independent and correlated activity of the HNH and RuvC domain. Cas9 was preloaded with sgRNA and incubated at 37 °C in 10-fold excess to 5 nM pM53.1 plasmid DNA containing the target sequence. For kinetic analysis, reactions were stopped by adding EDTA at different time points. Cas9 was digested by addition of 0.36 units of Proteinase K. The DNA fragments were resolved on a 0.9% agarose gel.

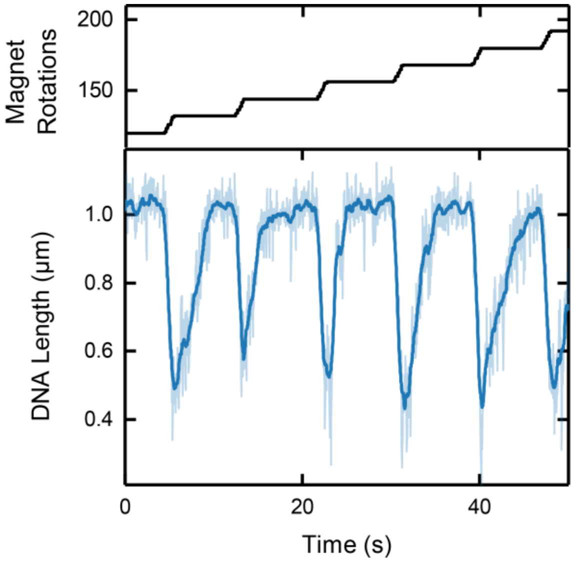

**Figure S2.** Characteristic magnetic tweezers trajectory for $Cas9_{wt}$ with repeated supercoiling of DNA. Loss of supercoiling indicates that the DNA has been nicked (non-target strand). Given that target strand cleavage precedes non-target strand cleavage (1), and the fact that this particular trace was recorded approximately two hours after nicking was first observed, we conclude that a double-strand break has been induced. Yet, the bead remains trackable, indicating that Cas9 remains bound and tightly holds on to the two DNA ends.

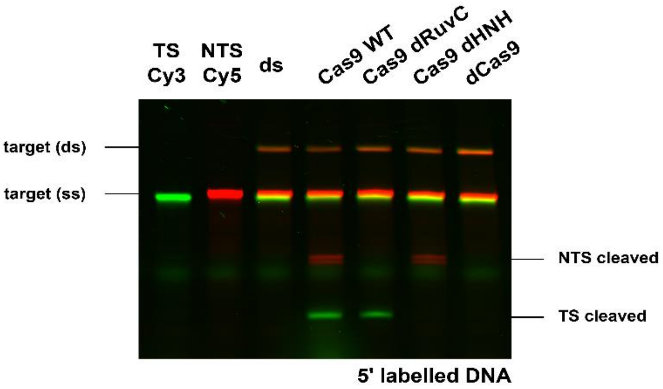

**Figure S3.** Fluorescently labeled DNA cleavage assay confirming the activity of $Cas9_{wt}$ and the nicking variants $Cas9_{dRuvC}$ and $Cas9_{dHNH}$. The target strand was labelled at the 5'-end with Cy3 (TS) and the non-target strand was 5' labeled with Cy5 (NTS). The double-stranded DNA was incubated for 60 minutes at 37°C with a 50-fold excess of a $Cas9_{X}$ preloaded with a crRNA:tracrRNA duplex. The reaction was stopped by addition of 0.36 units of Proteinase K. The samples were denatured at 95°C for five minutes and separated on a 15% denaturing polyacrylamide gel. Target (ds) indicates unseparated double-stranded DNA and target (ss) indicates denatured single strands.

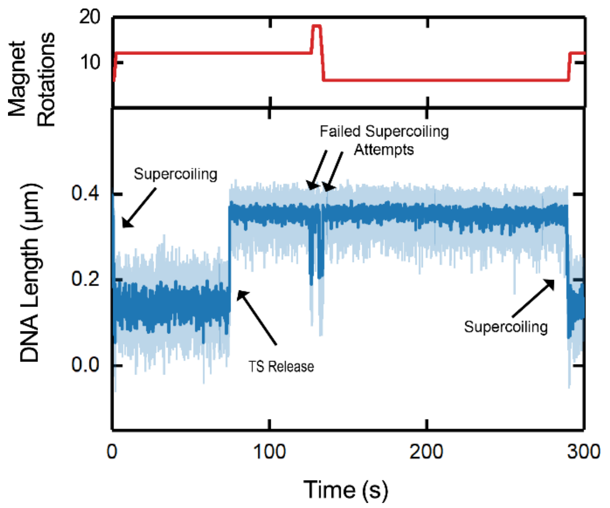

**Figure S4.** Characteristic magnetic tweezers trajectory of DNA length after nicking with $Cas9_{dRuvC}$ (light blue: Acquisition rate of 120 Hz; blue: smoothed to 6 Hz). Positive supercoiling is induced while Cas9 is bound to the target. After some time, the R-loop collapses and the supercoiling is lost due to the nick in the target strand. Subsequently, the nick in the DNA prevents further supercoiling until supposedly another Cas9 forms a new R-loop after approximately 100 s and the DNA regains its supercoilability.

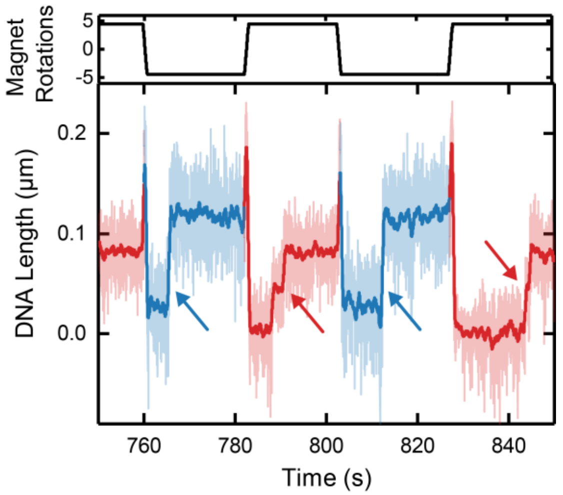

**Figure S5.** Characteristic magnetic tweezers trajectory for dCas9 (light blue/light red: acquisition rate of 120 Hz; blue/red: smoothed to 6Hz). R-loop formation (blue arrows) is performed at negative supercoiling (blue). R-loop collapse (red arrows) is performed at positive supercoiling (red). The process can be repeated multiple times when a new dCas9 protein binds to the DNA.

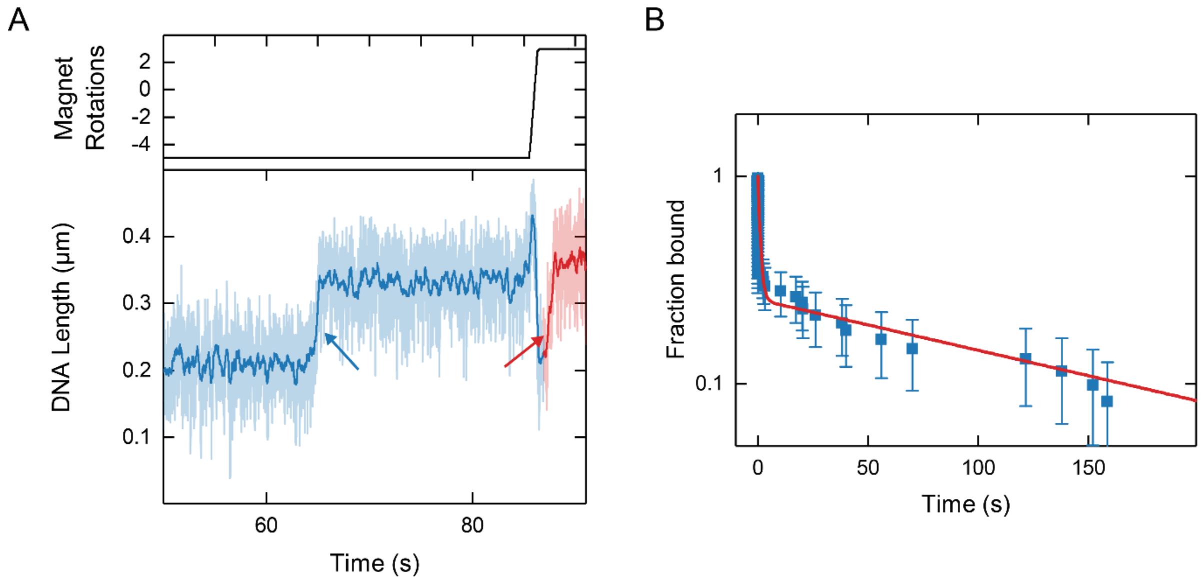

**Figure S6.** (A) Characteristic magnetic tweezers trajectory for dCas9 in $Mg^{2+}$-free buffer (light blue/light red: acquisition rate of 120 Hz; blue/red: smoothed to 6Hz). At negative supercoiling (blue), a full R-loop is formed (blue arrow), confirmed by the length difference of approximately 100 nm. At positive supercoiling (red), the R-loop collapses (red arrow), briefly visiting an intermediate R-loop state. (B) Kinetics of the R-loop collapse for dCas9 without magnesium (blue), inhibiting the conformational change into the docked state. A bi-exponential distribution is fitted to the data (red line). The average collapse time is 45 ± 33 s.

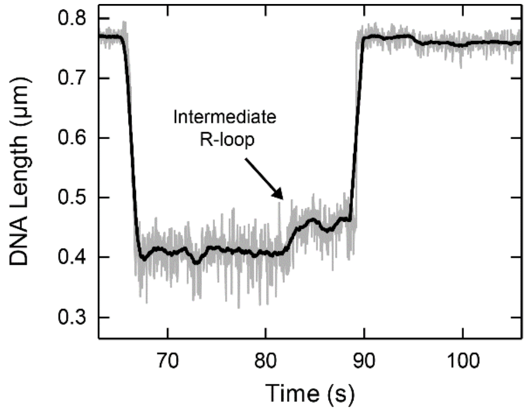

**Figure S7.** Characteristic magnetic tweezers trajectory for $Cas9_{dRuvC}$ dissociation in the unstable (‘undocked’) state. Supercoil release is preceded by an intermediate step, indicating that a full R-loop had been formed.

| Name | Sequence | Description |
| --- | --- | --- |
| sgRNA MT | **GAGCGAUACACAUGAGACGC**GUUUUAGAGCUAGAAAUAGCAAGUUAAAAUAAGGCUAGU CCGUUAUCAACUUGAAAAAGUGGCACCGAGUCGGUGCUUUUU | sgRNA for magnetic tweezers experiments |
| TK-386 | TAATACGACTCACTATAGAGCGATACACATGAGACGCGTTTTAGAGCTAGAAATAGC | |
| TK-626 | AAAAAGCACCGACTCGGTGCCACTTTTTCAAGTTGATAACGGACTAGCCTTATTTTAA CTTGCTATTTCTAGCTCTAAAAC | |
| FM | TS: GCTGACGTTTGTACTCCA**GCGTCTCATGTGTATCGCTC**AGCAGAGATTTCTGCT<br>NTS: AGCAGAAATCTCTGCT**GAGCGATACACATGAGACGCTGG**AGTACAAACGTCAGCT | 20nt protospacer fully matching the guide RNA |
| CRISPR-SpeI-For | GCGTAAGTCTCGAGAACTAGTTCCGTAAGATGCTTTTCTGTGACT | forward primer for all fragments |
| CRISPR-NotI-Rev-2_7 | GCGTAAGTGCGGCCGCTTCGTTCCACTGAGCGTCAGA | reverse primer for 2.7kbp fragment. |
| CRISPR-NotI-Rev-short | CGCTAAGTGCGGCCGCGCACAACATGGGGGATCATGTAACTCG | reverse primer for 1.5kbp fragment combined with CRISPR-SpeI-For |

**Table S1.** Sequences (5’ to 3’) of single-guide (sg) RNA, oligonucleotides used for sgRNA synthesis, dsDNA targets and PCR primers used for the magnetic tweezers experiments. The protospacers FM and T4 were inserted into a pUC19 plasmid pre-cleaved with SmaI. The PCR primers were used to amplify the DNA constructs. The protospacer, which is complementary to the guide RNA, is highlighted in blue. The PAM is highlighted in orange. The T7 promoter for RNA synthesis is highlighted in purple.

| Name | Sequence | Description |
| --- | --- | --- |
| sgRNA Bulk | **AUUGGUCACCUUACUUGGCA**GUUUUAGAGCUAGAAAUAGCAAGUUAAAAUAAGGCUA GUCCGUUAUCAACUUGAAAAAGUGGCACCGAGUCGGUGGUGCUUUUUUU | sgRNA for plasmid cleavage assay |
| Bulk Target | TS: **TGCCAAGTAAGGTGACCAAT**CCT<br>NTS: **AGGATTGGTCACCTTACTTGGCA** | target and non-target strand for plasmid cleavage assay |
| crRNA | CGCGCCGAGGUGAAGUUCGAGUUUUAGAGCUAUGCUGUUUUG | crRNA for oligomer fluorescence cleavage assay |
| tracrRNA | GGACAGCAUAGCAAGUUAAAAUAAGGCUAGUCCGUUAUCAACUU GAAAAAGUGGCACCGAGUCGGUGCUUUUU | tracrRNA for oligomer fluorescence cleavage assay |
| Fluorescence Target | TS: ACCAG**GGTGTCGCCCTCGAACTTCA**CCTCGGCGCGGGTCTTGTA<br>NTS: TACAAGACCCGCGCCG**AGGTGAAGTTCGAGGGCGACACC**CTGGT | target and non-target strand for fluorescence oligomer cleavage assay |

**Table S2.** Protospacers and PCR primers (5’ to 3’) used in bulk cleavage assays.
